# Biological Foundation Models Enable CRISPR Array Detection Without Metagenomic Assembly

**DOI:** 10.64898/2026.02.16.706169

**Authors:** Lukas Daugaard Schröder, Ryan Köksal, Alexander Mitrofanov, Michael Uhl, Rolf Backofen

**Affiliations:** Chair of Bioinformatics, University of Freiburg, Georges-Köhler-Allee 101, 79110, Freiburg, Germany; Signalling Research Centres BIOSS and CIBSS, University of Freiburg, 79085, Freiburg, Germany

**Keywords:** CRISPR, CRISPR-Cas, Machine learning, Deep learning, Foundation models, Metagenomics

## Abstract

Accurate identification of CRISPR arrays is essential for studying prokaryotic adaptive immunity, yet existing tools struggle with short-read sequencing data and arrays containing degenerate repeats. These limitations restrict CRISPR analysis in metagenomic and fragmented genomic datasets. We present a foundation model-based approach for CRISPR array detection that addresses both these challenges. We fine-tune a large genomic foundation model using the Parameter-Efficient Fine-Tuning (PEFT) method, Low-Rank Adaptation (LoRA) to perform per-nucleotide classification of DNA sequences into repeat, spacer, and non-array regions directly from raw input nucleotide sequences. We develop two model variants for different sequence context lengths. The long-context model supporting sequences of up to 8,192 nucleotides achieves 98.16% test accuracy and identifies degenerate repeat candidates missed by similarity-based CRISPR detection tools. The short-context model supports sequences of up to 150 nucleotides, optimized for Illumina reads, reaches 90.03% accuracy and enables direct analysis of individual reads without assembly. On simulated metagenomic data, it achieves a spacer recall of 49.12% and recovers 12.57% of spacers that are otherwise not detected by dedicated metagenomic CRISPR array detection methods which require metagenomic assembly. Together, these results demonstrate that genomic foundation models provide a robust and complementary paradigm for CRISPR array detection.

## 1 Introduction

CRISPR arrays are a defining component of CRISPR-Cas adaptive immune systems in bacteria and archaea, providing a molecular record of past infections. The accurate identification of these arrays is central to studying microbial immunity, CRISPR-Cas diversity, and host-virus coevolution [1–4].

Early CRISPR detection tools such as CRT [5] and PILER-CR [6] identify regularly spaced, highly similar direct repeats and were developed primarily for complete or near-complete genomes. More recent approaches, including CRISPRCasFinder [7] and CRISPRidentify [8], improve filtering, confidence estimation, and sensitivity through refined heuristics and machine learning. Despite these advances, existing methods largely assume long contiguous sequences and multiple repeat instances, limiting their robustness under sequence fragmentation or repeat degeneration.

These assumptions are frequently violated in metagenomic sequencing, where data consists of short reads or highly fragmented contiguous sequences. In this setting, CRISPR arrays are often truncated, split across reads, or reduced to single repeat-spacer units, and assembly-based workflows may discard CRISPR loci during graph simplification. As a result, both genome-oriented and metagenomic approaches suffer from reduced sensitivity, although for different reasons. We hypothesize that these failures arise from shared reliance on explicit repeat detection and rigid structural criteria rather than from individual algorithmic choices.

To address these limitations, we reformulate CRISPR array detection as a context-aware, per-nucleotide sequence labeling problem using deep genomic foundation models. By operating directly on sequence context and integrating information across variable-length inputs, this approach does not require prior repeat enumeration or metagenomic assembly and remains effective on fragmented or degenerated arrays. In the following sections, we show that this strategy enables robust identification of CRISPR array regions in both complete genomes and short-read–derived sequences, including cases where classical methods fail.

## 2 Methods

### 2.1 Data

To fine-tune Evo on this task, we obtained high-confidence CRISPR array annotations by running CRISPRidentify [8] on a publicly available genomic dataset, comprising of 47,760 complete prokaryotic genomes from [9–11]. We retained only arrays classified as *bona fide* with confidence scores of at least 0.75. This ensures high annotation quality is available for fine-tuning to complement the large amount of lower quality data used when pre-training Evo. For each array, we included up to 500 nucleotides of flanking genomic sequence on each side of the array interval where available. We labeled these flanking regions as non-array to enable the model to learn CRISPR array boundaries.

Because the dataset often contains different strains of the same organism, e.g., *Escherichia coli*, the identified CRISPR arrays are frequently identical or nearly identical, differing by as little as a single spacer. The presence of such highly similar arrays in both the training and test sets would introduce data leakage and lead to overly optimistic performance estimates. To ensure a valid and unbiased evaluation, we therefore performed de-duplication prior to dataset splitting. This yielded a final dataset of 5,084 unique CRISPR arrays, with a median CRISPR array length of 701 nucleotides. The shortest array in that dataset had 143 nucleotides, while the longest reached 26,001, reflecting substantial length variability within the dataset (Refer to Figure S1 for details). Repeat and spacer nucleotide classes were approximately balanced, while non-array nucleotides constitute the largest class in the dataset. Detailed dataset statistics are provided in Supplementary Figure S2.

We split the dataset at the sequence level using a 70-10-20 split for training, validation, and test sets, respectively, ensuring no leakage of sequences between splits.

### 2.2 Fine-tuning for CRISPR Array Detection

We formulate CRISPR array detection as a multi-class per-nucleotide sequence labeling task with three classes assigned to each nucleotide: repeat, spacer, and non-array. We used Evo [12], a genomic foundation model pretrained on whole prokaryotic genomes with a total of 300 billion nucleotides. Specifically, we fine-tune the Evo-1-8k-base variant, which supports context lengths of up to 8,192 nucleotides and contains 7 billion parameters. Evo uses a byte-level tokenization scheme, where nucleotides (A, C, G, T) are encoded directly as UTF-8 bytes.

To adapt the pretrained Evo model to the CRISPR labeling task, while preserving its learned genomic knowledge and minimizing computational requirements, we employed Low-Rank Adaptation (LoRA) [13]. LoRA introduces trainable low-rank decomposition matrices into Evo’s query-key-value projection in the attention layers and its three linear layers without modifying the original pretrained weights. This allowed us to fine-tune the Evo backbone on our high-confidence annotations while updating a small fraction of Evo’s total parameters. This enables the model to learn specific CRISPR representations from high quality annotations while still retaining Evo’s pretrained general genomic knowledge. We apply LoRA with a rank **r** of 1 and a scaling factor *α* of 32. We additionally employ dropout of 0.05 to LoRA layers. Details of the hyperparameter search space and the final configuration used are provided in Supplementary Table 1.

We added a task-specific classification head that maps the final hidden representations of the model to class logits for the three output classes (repeat, spacer, non-array).

For fine-tuning, we employed the AdamW optimizer [14], with decoupled weight decay and a per-nucleotide cross-entropy loss. We additionally implemented learning-rate warmup, followed by decay and gradient clipping. Due to memory constraints, we used a batch size of 1 and did not employ gradient accumulation as we found it to worsen accuracy empirically. A schematic overview of the approach is shown in Figure 1.

**Fig. 1.**
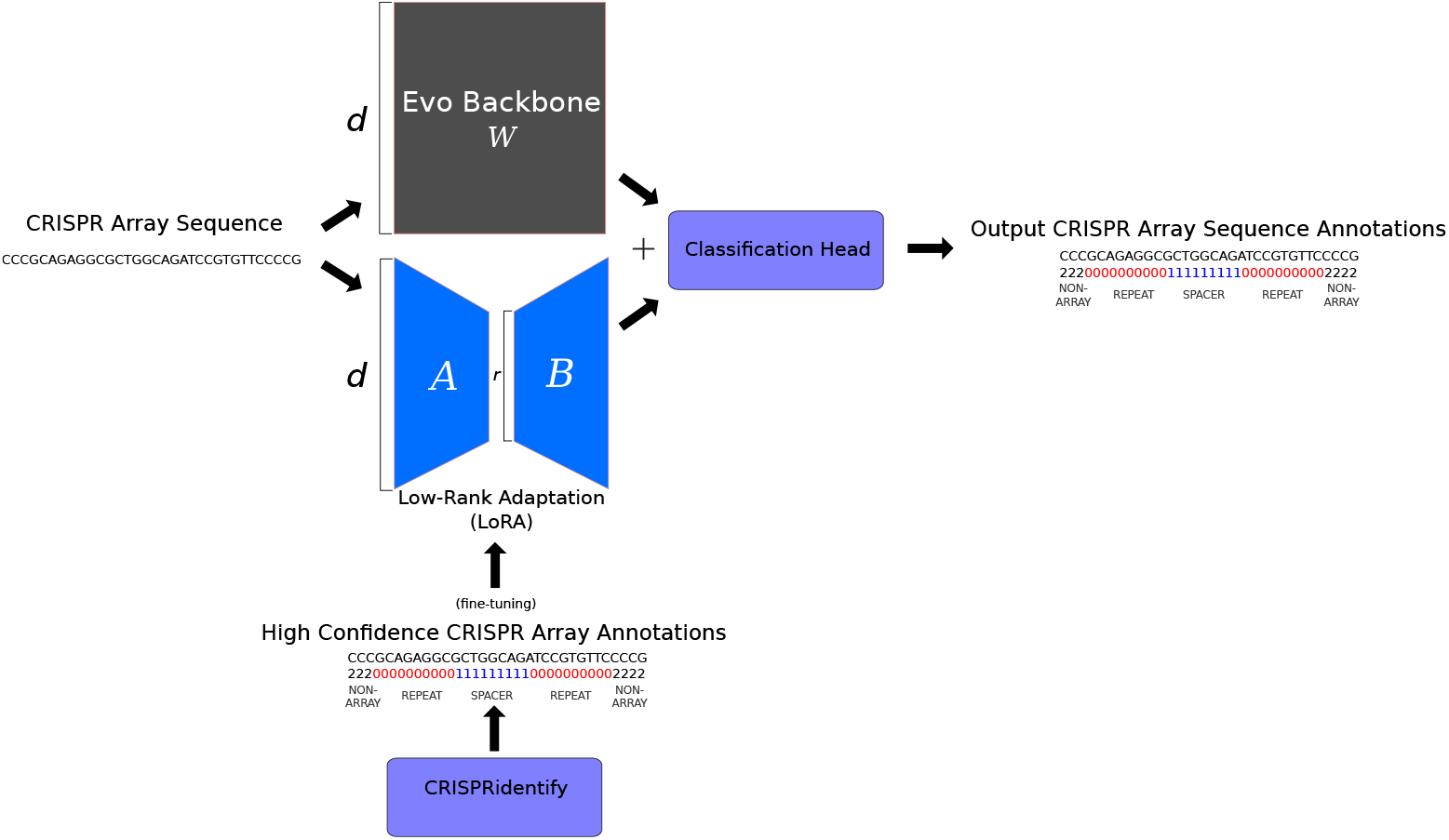
We fine-tuned the pretrained genomic foundation model, Evo, originally optimized for next-nucleotide prediction, to perform a per-nucleotide classification task. Using CRISPRidentify, we obtained high-confidence CRISPR array annotations, with each nucleotide in each sequence labeled as repeat, spacer or non-array. We then fine-tune Evo using Low-rank Adaptation (LoRA) to learn CRISPR representations from the high confidence sequences while retaining general prokaryotic knowledge from pre-training.

We selected hyperparameters using grid search optimized on the validation set described in Section 2.1. We conducted a separate hyperparameter optimization for long-context (8,192 nt) and short-context (150 nt) models due to differences in effective batch size and memory constraints, following practices recommended by [15–17]. Details of the range of hyperparameter values considered in this search and the final configuration are provided in Supplementary Table 1.

All models were implemented using PyTorch [18] and the Hugging Face Transformers library [19]. All experiments were performed on a single NVIDIA H200 141GB GPU.

### 2.3 Evaluation Metrics

We evaluated model performance primarily using per-nucleotide classification accuracy on held-out test sequences. To better characterize error modes, we additionally analyzed class-specific mis-classification accuracy, with particular attention to errors at CRISPR array boundaries and within degenerated repeat regions.

For metagenomic evaluation, we extracted predicted spacer sequences from model outputs and compared them against validated spacer collections from CRISPRCasDB [20]. Spacer recall was defined as the fraction of validated spacers recovered by the model.

To assess the detection of degenerated repeat regions, we analyzed predicted repeat segments extending beyond annotated array boundaries. These segments were aligned to the corresponding array consensus repeats and alignments exceeding a significance threshold were counted as supporting evidence for degenerate repeat recovery.

## 3 Results

### 3.1 Zero-Shot Next-Nucleotide Analysis

We first evaluated the behavior of the pretrained Evo foundation model in a zero-shot setting by analyzing raw next-nucleotide prediction probabilities without any task-specific fine-tuning. Across genomic regions containing annotated CRISPR arrays, the model achieved an average next-nucleotide prediction probability of 57.22% on the fine-tuning dataset.

We observed that high-confidence predictions were strongly enriched in CRISPR repeat regions. Among nucleotides with above-average prediction probability, 75.00% corresponded to repeats, compared to 21.11% non-array regions and 3.89% spacers. Figure 2 shows a representative example of next-nucleotide prediction probabilities across an annotated CRISPR array, illustrating consistently elevated confidence within repeat regions relative to spacers and surrounding genomic background. Within repeats, prediction probabilities frequently exceeded 95% for most nucleotide positions, while reduced confidence was observed at array boundaries and within spacer regions.

**Fig. 2.**
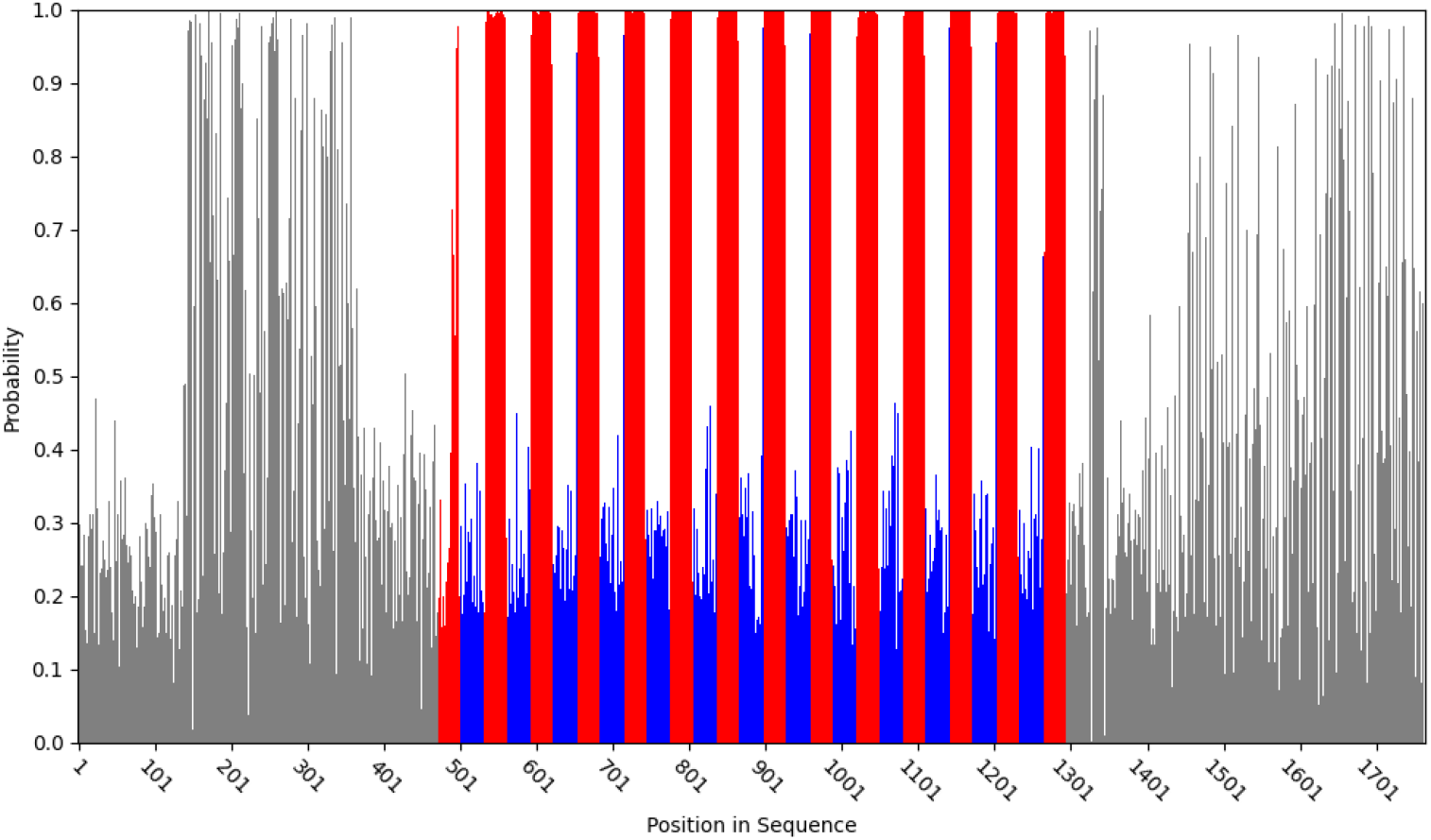
Zero-shot next-nucleotide prediction confidence highlights CRISPR repeat structure by Evo. Next-nucleotide prediction probabilities from the pretrained Evo model are shown across a genomic region containing a CRISPR array detected by CRISPRidentify (genome accession number APZF01000100, positions 167,361– 168,247). Nucleotides are colored by CRISPRidentify annotation, with repeats shown in red, spacers in blue, and non-array regions in grey. Prediction confidence closely matches the annotated intervals, with consistently elevated values in repeat regions and reduced values in spacers and flanking genomic background and pronounced drops at array boundaries. This alignment is consistent with the conserved and structurally constrained nature of CRISPR repeats and suggests that Evo’s pretrained representations encode CRISPR-associated sequence structure even without CRISPR-specific supervision.

These observations indicate that the pretrained model captures sequence regularities characteristic of CRISPR arrays, particularly repeat-associated structure, despite the absence of CRISPR-specific supervision. However, this analysis reflects probabilistic sequence modeling behavior rather than explicit CRISPR array detection.

Given this level of pretrained performance, even a simple threshold-based binary classifier that labels positions with a predicted probability greater than 90% as repeat nucleotides and all others as spacer nucleotide can accurately partition CRISPR array regions into repeats and spacers, achieving an average classification accuracy of 81%.

### 3.2 Supervised Fine-Tuning Improves Per-Nucleotide CRISPR Classification

We next fine-tuned the model using per-nucleotide CRISPR array annotations and evaluated performance on held-out genomic regions. Fine-tuning substantially improved discrimination between CRISPR array components and surrounding genomic background, producing spatially coherent predictions across repeat and spacer regions.

Accurately distinguishing between repeats, spacers, and non-array regions is an essential part of CRISPR array detection. Compared to the zero-shot setting, the fine-tuned model showed a marked improvement in distinguishing between spacers and non-array regions while maintaining high sensitivity within annotated arrays. Residual errors were predominantly localized to array boundaries and regions with highly degenerated repeats (See Supplementary Figure S3 for details). Predictions remained stable under realistic input window shifts and sequence fragmentation, supporting applicability in diverse sequencing contexts.

### 3.3 Accuracy Across Input Lengths

To assess robustness under varying sequence contiguity, we evaluated per-nucleotide classification accuracy across different target lengths. Figure 3 summarizes test accuracy for models trained with 150 nt and 8,192 nt contexts. We found that model performance depends on the amount of available sequence context. Shorter target lengths resulted in a moderate decrease in classification accuracy, whereas longer input sequences consistently improved performance. This pattern indicates that additional contextual information facilitates more accurate identification of repeat–spacer structure and CRISPR array boundaries.

**Fig. 3.**
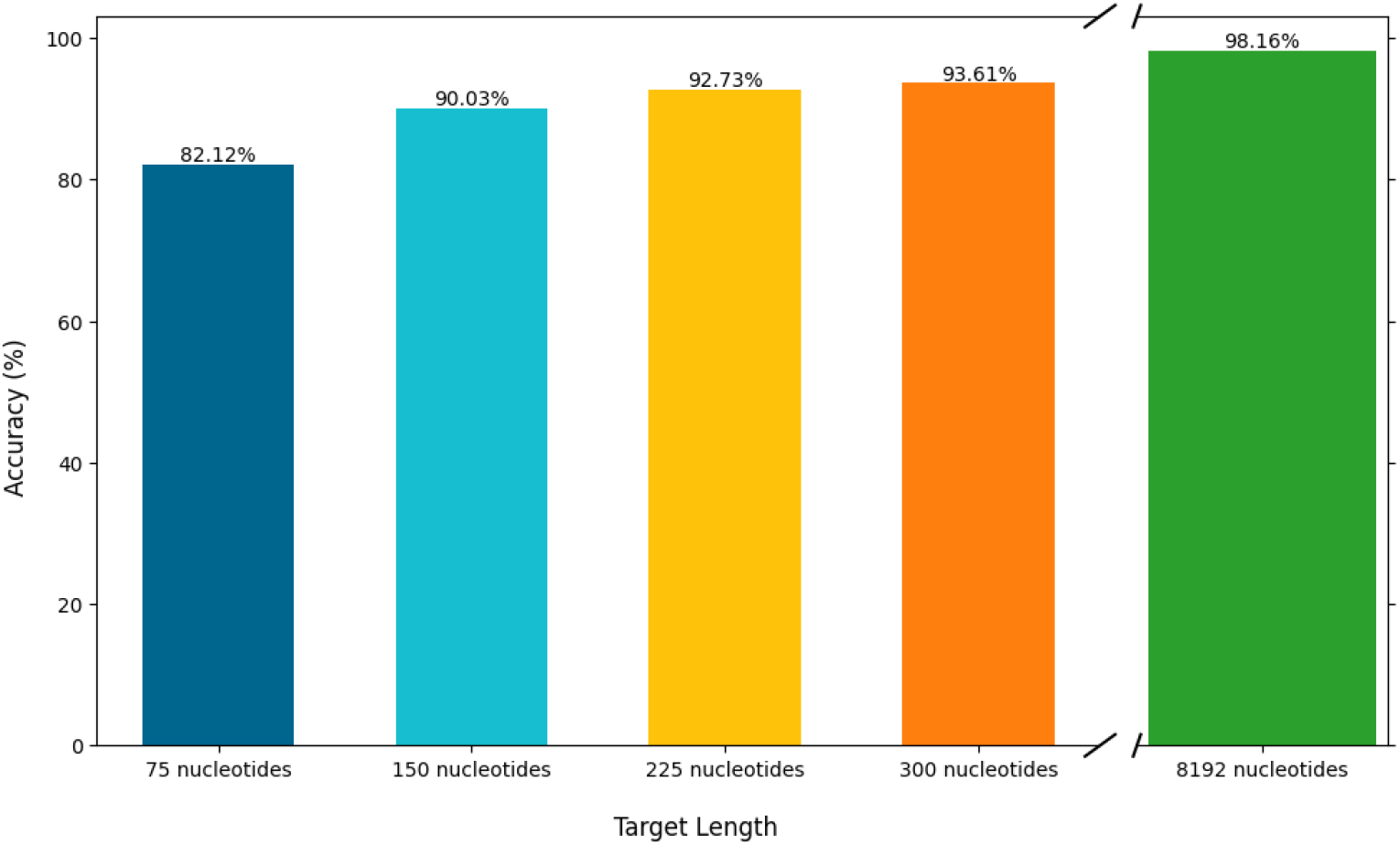
Per-nucleotide classification accuracy of Evo models fine-tuned to label repeat, spacer, and non-array regions is shown for different input sequence lengths. Model accuracy increases with available sequence context, indicating that longer-range information improves resolution of CRISPR array structure and boundaries. Nevertheless, the short-context model (150 nt) achieves high accuracy of 90.03%, demonstrating that CRISPR array components can be reliably classified even in highly fragmented sequences such as individual short sequencing reads.

Despite reduced context, we observed stable performance across a broad range of input lengths. This demonstrates that the model does not rely exclusively on long-range dependencies but instead leverages local sequence features captured during pretraining and reinforced through fine-tuning.

Overall, we show that the approach generalizes effectively across different sequencing regimes, supporting its applicability to both long assembled genomic sequences and short-read or metagenomic data where input contiguity is inherently limited.

### 3.4 Short-Read Metagenomic Spacer Recovery

We evaluated the fine-tuned short-context (150 nt) model on simulated short-read metagenomic data and compared spacer recovery to MCAAT [21]. Unlike assembly-based approaches, our method operates directly on individual sequencing reads and does not require contig reconstruction or graph-based pre-processing. This enables CRISPR signal detection in highly fragmented metagenomic datasets, where assembly may be incomplete or fail entirely.

In order to quantify the findings, we compute a recall metric by measuring how many spacers in the CRISPRCasdb [20] are recovered in our dataset. Spacers from the two sets are compared using BLAST [22], with a minimum percentage identity of 95%, a minimum fraction of the query aligned of 90%, and an E-value threshold of 10^−5^, so that only highly significant matches are considered.

The model achieved a spacer recall of 49.12% and recovered complementary signal, with 12.57% of validated spacers detected exclusively by our approach (Figure 4). This demonstrates that the foundation model identifies biologically meaningful spacers that are missed by methods relying on assembly-derived contigs.

**Fig. 4.**
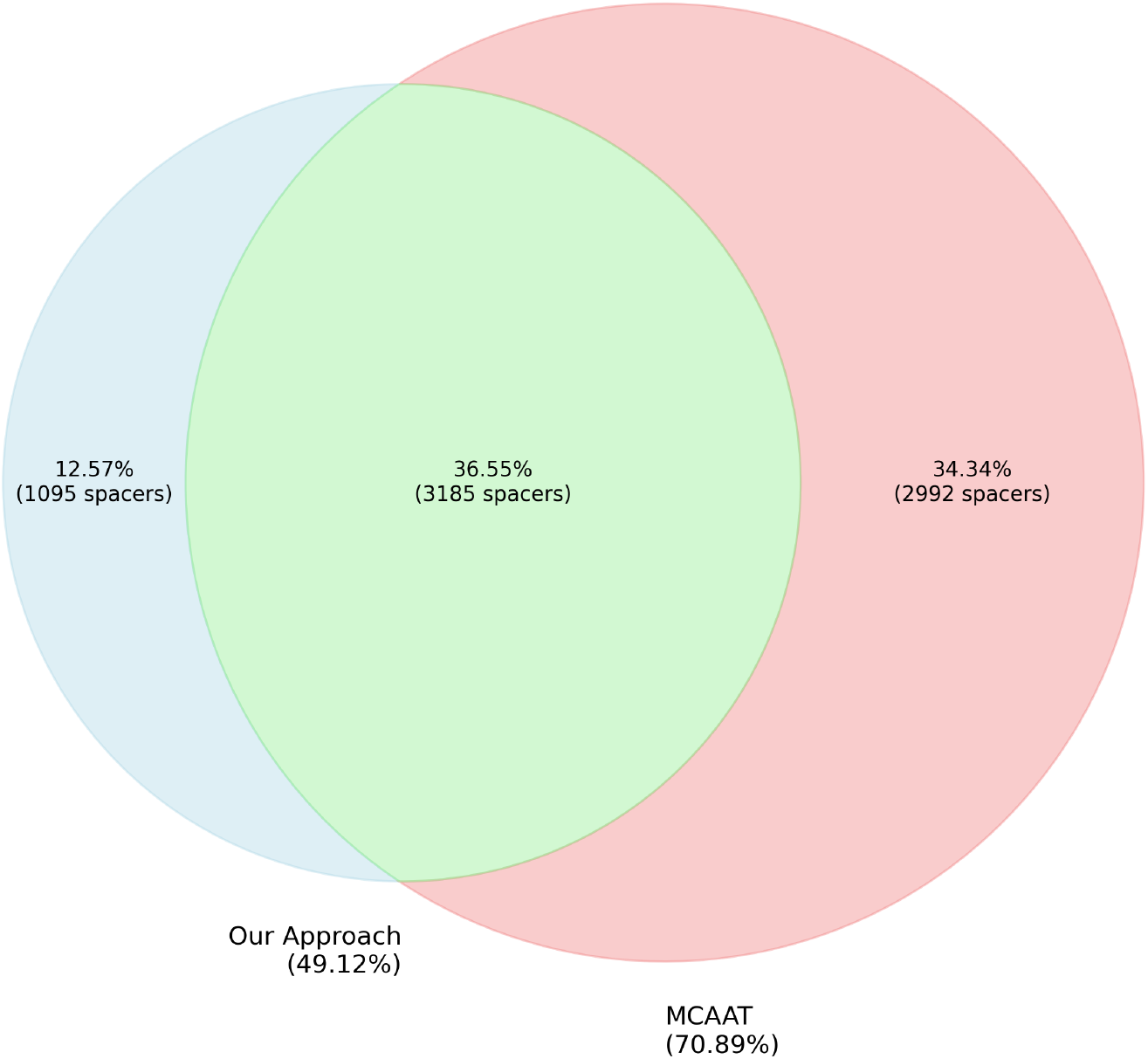
Overlap of validated CRISPR spacers recovered from simulated metagenomic short-read data by the short-context (150 nt) model variant and MCAAT. While MCAAT achieves higher overall spacer recall (70.89 %), our fine-tuned Evo model recovers a distinct subset of validated spacers directly from individual reads without assembly, achieving a recall of 49.12 %. Notably, 12.57 % of spacers detected by our approach are not recovered by MCAAT. This demonstrates that combining both approaches increases the total number of recovered spacers and highlights their complementary strengths for metagenomic CRISPR analysis.

Importantly, the short-context model is effective on reads as short as 150 nucleotides, a length at which only a single repeat or spacer may be present. This capability is particularly relevant for Illumina-based metagenomic datasets, where canonical CRISPR array structure is rarely fully contained within a single fragment. Moreover, by leveraging learned sequence context rather than exact repeat matching, the model remains sensitive to CRISPR repeats that harbor mutations. Such mutations can disrupt k-mer continuity and cause De Bruijn graph–based assembly methods to fragment or discard CRISPR loci. The ability of our model to recover spacers under these conditions highlights the biological utility of an assembly-free, context-aware approach for studying CRISPR systems in evolving microbial communities.

### 3.5 Detection of Degenerate Repeat Regions

Finally, we examined whether the fine-tuned model identifies degenerated repeat regions extending beyond annotated CRISPR array boundaries. We detected 71 candidate regions with predicted CRISPR signal outside annotated arrays, of which 92.5% aligned significantly to the corresponding array consensus repeats (Refer to Supplementary Tables 2-3 for details). These regions likely represent truncated or degenerated repeat elements that are difficult to detect using explicit repeat-based criteria.

## 4 Discussion

Our results demonstrate that genomic foundation models offer a biologically grounded alternative to explicit repeat-based CRISPR detection. Even without task-specific supervision, the pretrained model captures conserved sequence patterns characteristic of CRISPR repeats, while supervised fine-tuning enables accurate per-nucleotide discrimination of repeats, spacers, and background sequence across variable input lengths.

A key advantage of this approach is its ability to operate directly on short sequencing reads without assembly. This is particularly important in metagenomic datasets, where CRISPR arrays are frequently fragmented or affected by sequence variation, causing assembly- and similarity-based methods to miss or discard CRISPR loci. Furthermore, by modeling sequence context rather than exact repeat identity, the foundation model remains sensitive to degenerated and mutated repeats, enabling recovery of biologically meaningful CRISPR elements in diverse and evolving microbial communities.

## Funding

This research was funded by the Deutsche Forschungsgemeinschaft (DFG, German Research Foundation) under grant number 539134284, through EFRE (FEIH 2698644) and the state of Baden-Württemberg; Deutsche Forschungsgemeinschaft (DFG, German Research Foundation) [BA 2168/23-1/2 SPP 2141]; Much more than Defence: the Multiple Functions and Facets of CRISPR–Cas; Funding for open access charge. Probabilistic Structures in Evolution [BA 2168/23-1/2 SPP 2141]; and the state of Baden-Württemberg through bwHPC and the German Research Foundation (DFG) [grant number INST 35/1597-1 FUGG].

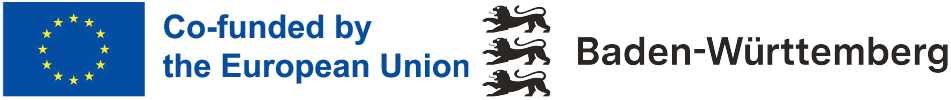

## Data availability

All fine-tuned models, code, and data required to reproduce the results in this publication are publicly available at https://github.com/ivelet/CRISPREvo.

## 6 Author contributions statement

L.D.S. implemented the model and training procedures, hypothesized the use of Evo, conducted fine-tuning experiments, performed computational analyses, contributed to discussions, and revised the manuscript. R.K. designed and performed fine-tuning experiments, developed the model architecture and training procedures, supervised the study, contributed to discussions, and wrote the manuscript. A.M. conceptualized the study, designed and performed CRISPR-related experiments and analyses, prepared all CRISPR datasets, supervised the study, contributed to discussions, and wrote the manuscript. M.U. designed experiments, contributed to discussions and interpretation of results, performed analytical review, and wrote the manuscript. R.B. supervised the study, provided funding, contributed to discussions, and revised the manuscript.

## Supplementary Materials

### S1. Dataset Composition

Distribution of nucleotide classes and CRISPR array lengths in the training dataset.

**Fig. S1.**
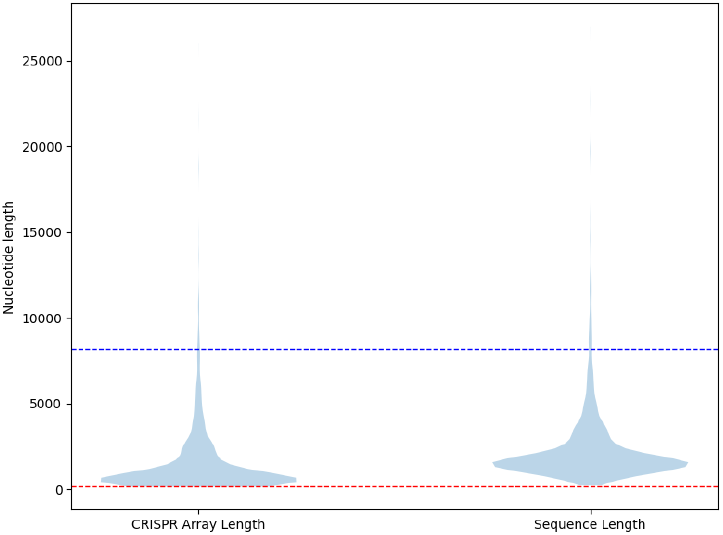
Distribution of Finetune Dataset Lengths. Lengths of the 5,084 CRISPR arrays and the length of their sequences (CRISPR array length plus flanking nucleotides) in the fine-tuning dataset. The violin plot shows the distribution of lengths. The median lengths are 701 and 1,639, the longest are 26,001 and 27,001, and the shortest are 143 and 221, for CRISPR array length and sequence length, respectively. The red dashed line represents the target length of the short-context model (150 nt) and the blue line is of the long-context model (8,192 nt). This distribution shows that the majority of CRISPR arrays fit within the 8,192 context window and that all sequences are longer than the 150 target length.

**Fig. S2.**
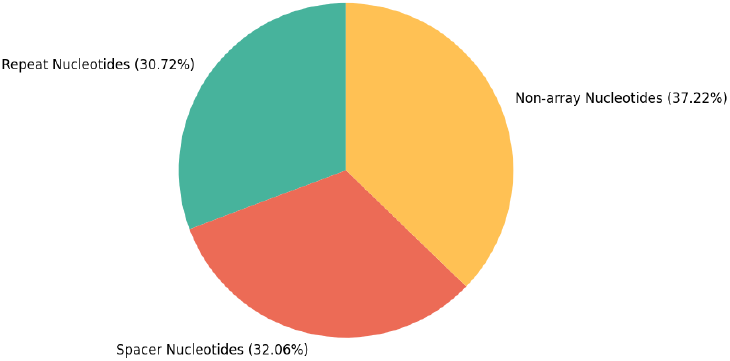
Nucleotide Distribution by Class of the fine-tuning Dataset. The chart shows the proportion of nucleotides by each label: repeat nucleotides (3,563,841), spacer nucleotides (3,719,636), and non-CRISPR array (non-array) nucleotides (4,318,435). This distribution shows that repeat and spacer nucleotide classes are balanced, while non-array nucleotides form the largest class.

### S2. Error-Type Breakdown Across Input Lengths

Class-specific error distributions for short- and long-context models.

**Fig. S3.**
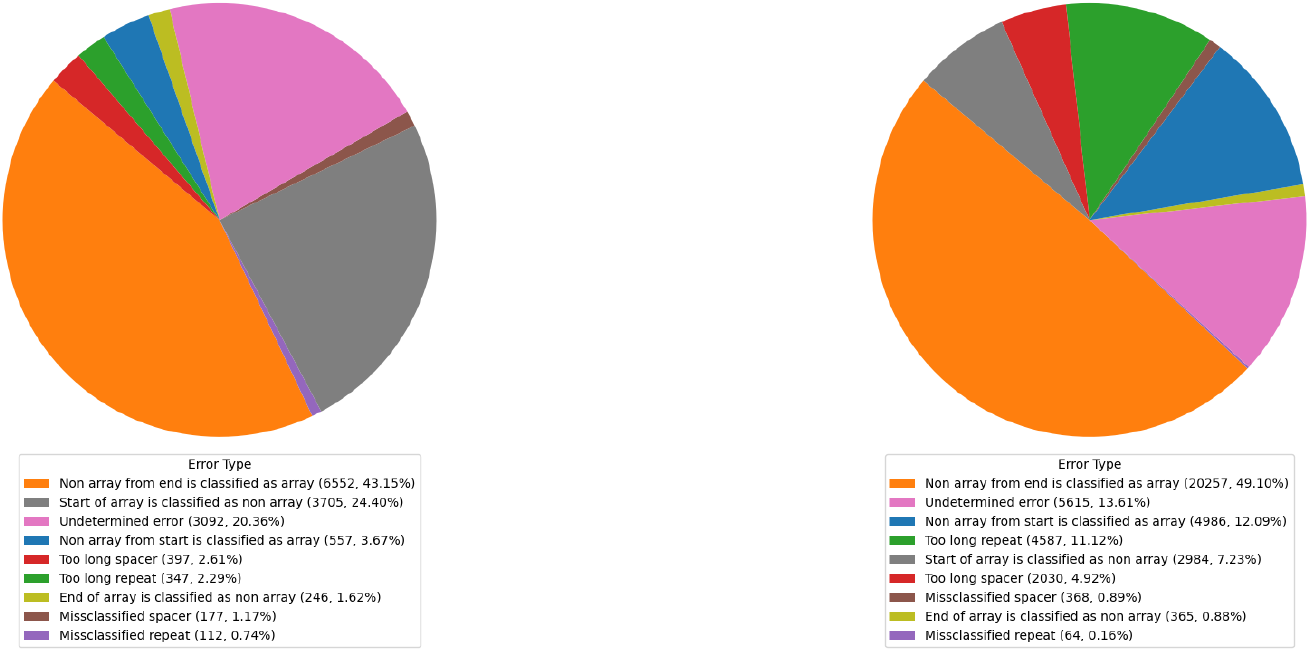
Error-type breakdown for the 150-nt model (top) and the 8,192-nt model (bottom).

### S3. Hyperparameter Optimization

**Table 1.** We performed grid search over a given range of values to select the optimal hyperparameters for fine-tuning.

| Hyperparameter | Search Range | Selected Value |
| --- | --- | --- |
| Learning rate | $\{1 \times 10^{-5}, 1 \times 10^{-3}\}$ | $1 \times 10^{-4}$ |
| Weight decay | $\{0, 1 \times 10^{-4}, 1 \times 10^{-3}\}$ | 0 |
| Batch size | $\{1, 2, 4, 8, 16\}$ | 2 |
| Dropout | $\{0.0, 0.1, 0.2\}$ | 0.10 |
| Warmup steps | $\{0, 500, 1000\}$ | 0 |
| Training epochs | $\{1, 10\}$ | 3 |
| LoRA rank | $\{4, 8, 16\}$ | 1 |
| LoRA $\alpha$ | $\{4, 8, 16, 32\}$ | 32 |
| LoRA dropout | $\{0, 0.05, 0.10, 0.15, 0.20\}$ | 0.05 |

### S4. Degenerate repeat candidates

**Table 2.**
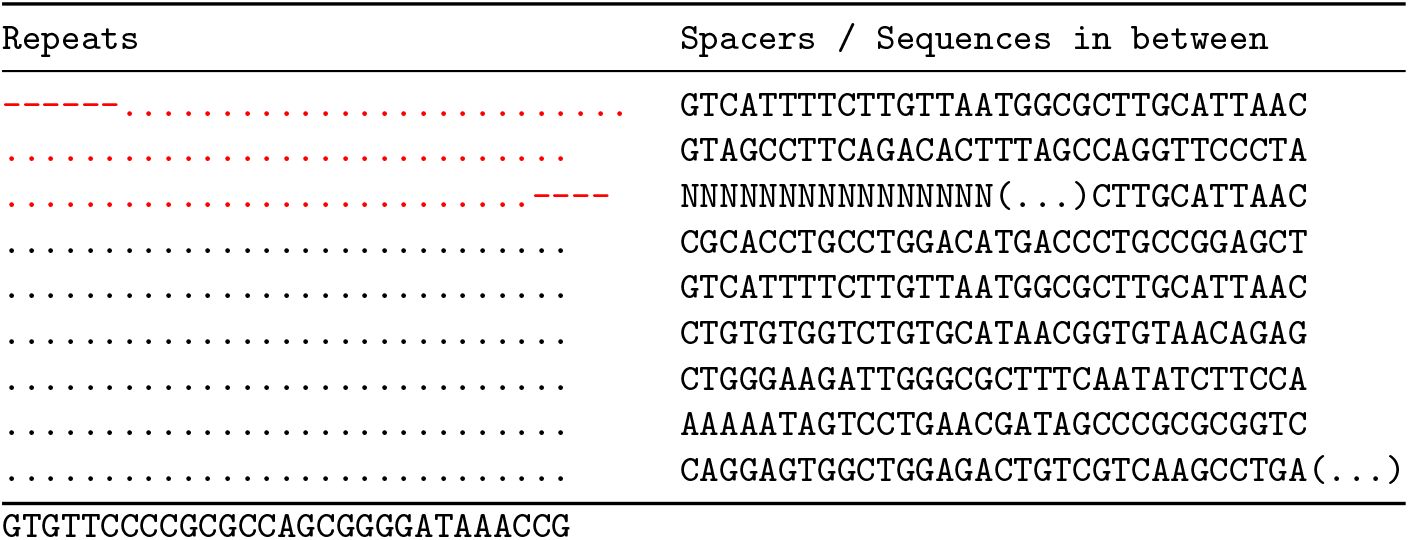
An example of a CRISPR array detected in accession CASA01000002 (positions 108,834–109,415), illustrating repeat and spacer organization. Repeats are shown on the left and spacers or intervening sequences on the right. Differences from the consensus repeat (as defined by CRISPRidentify) are indicated in red to mark a candidate degenerated repeat. Only nucleotide positions that deviate from the consensus are displayed for clarity.

**Table 3.**
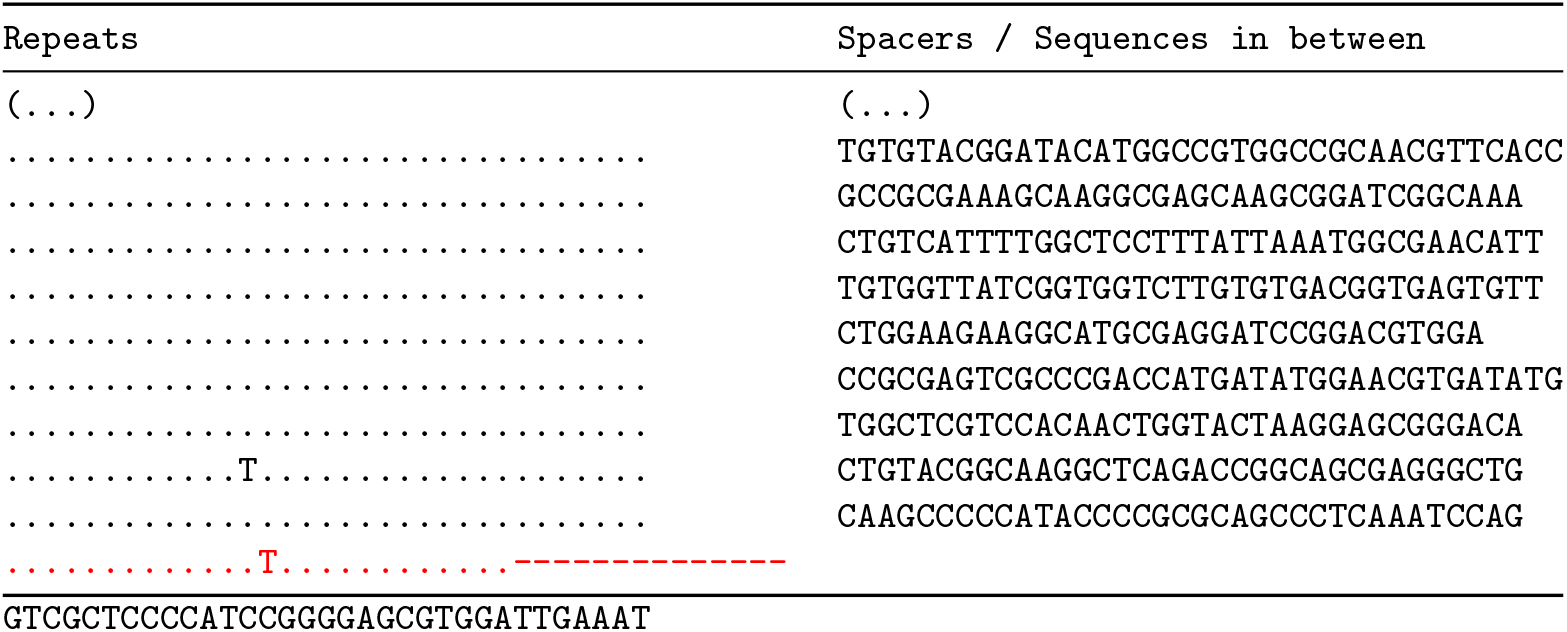
An example of a CRISPR array detected in accession CP034192 (positions 1,672,883–1,674,884), illustrating repeat and spacer organization. Repeats are shown on the left and spacers or intervening sequences on the right. Differences from the consensus repeat (as defined by CRISPRidentify) are indicated in red to mark a candidate degenerated repeat. Only nucleotide positions that deviate from the consensus are displayed for clarity.

